# SARS-CoV-2 susceptibility of cell lines and substrates commonly used in diagnosis and isolation of influenza and other viruses

**DOI:** 10.1101/2021.01.04.425336

**Authors:** Li Wang, Xiaoyu Fan, Gaston Bonenfant, Dan Cui, Jaber Hossain, Nannan Jiang, Gloria Larson, Michael Currier, Jimma Liddell, Malania Wilson, Azaibi Tamin, Jennifer Harcourt, Jessica Ciomperlik-Patton, Hong Pang, Naomi Dybdahl-Sissoko, Ray Campagnoli, Pei-Yong Shi, John Barnes, Natalie J. Thornburg, David E. Wentworth, Bin Zhou

**Affiliations:** Centers for Disease Control and Prevention, Atlanta, Georgia, USA; Oak Ridge Institute for Science and Education, Oak Ridge, Tennessee, USA; Battelle Memorial Institute, Atlanta, Georgia, USA; University of Texas Medical Branch, Galveston, Texas, USA

**Author notes:** Correspondence: Bin Zhou.

## Abstract

Coinfection with severe acute respiratory syndrome coronavirus 2 (SARS-CoV-2) and other viruses is inevitable as the COVID-19 pandemic continues. This study aimed to evaluate cell lines commonly used in virus diagnosis and isolation for their susceptibility to SARS-CoV-2. While multiple kidney cell lines from monkeys were susceptible and permissive to SARS-CoV-2, many cell types derived from human, dog, mink, cat, mouse, or chicken were not. Analysis of MDCK cells, which are most commonly used for surveillance and study of influenza viruses, demonstrated that they were insusceptible to SARS-CoV-2 and that the cellular barrier to productive infection was due to low expression level of the angiotensin converting enzyme 2 (ACE2) receptor and lower receptor affinity to SARS-CoV-2 spike, which could be overcome by over-expression of canine ACE2 in trans. Moreover, SARS-CoV-2 cell tropism did not appear to be affected by a D614G mutation in the spike protein.

## INTRODUCTION

Coronavirus Disease 2019 (COVID-19) has resulted in more than 70 million laboratory confirmed cases and more than 1.6 million deaths in less than a year since the first case was confirmed. Coinfection with SARS-CoV-2 and other viruses, such as influenza virus, has been reported (1–4). As cases of COVID-19 continue to climb sharply, more coinfections are expected, especially in the current and future influenza seasons.

Isolation and propagation of virus from clinical specimens in cell cultures or embryonated chicken eggs are widely used for virus diagnosis and vaccine production, mostly under biosafety level 2 (BSL2) containment. Currently, SARS-CoV-2 should only be isolated and propagated under BSL3 containment due to its risk to laboratorians and the general public. Therefore, if any of these cell lines or eggs support productive replication of SARS-CoV-2, then a validated procedure should be implemented to rule out the presence of SARS-CoV-2 in the specimens prior to their inoculation. However, adding a diagnostic step specific to SARS-CoV-2 in many circumstances is impractical or substantially increases the cost and labor required.

We conducted the present study to determine whether cell lines and eggs commonly used for isolation and propagation of influenza virus, poliovirus and other human viruses can support productive replication of SARS-CoV-2. If a substrate is confirmed to be insusceptible to SARS-CoV-2, modification of procedures for diagnosis and isolation of susceptible viruses in that substrate may be unnecessary. While all results were repeated under the same or slightly different conditions, some of our results were further confirmed using two divergent SARS-CoV-2 strains, with multiple assay methods, and in cell lines from different sources.

Our study provides important information on the risk of inadvertent propagation of SARS-CoV-2 in cell lines and/or substrates when conducting diagnosis, isolation, propagation, or vaccine production of other viruses.

## MATERIALS AND METHODS

### Viruses

SARS-CoV-2/USA-WA1/2020 (USA-WA1) was isolated from the specimen of the first confirmed case in the United States as described previously (5). SARS-CoV-2/Massachusetts/VPT1/2020 (MA/VPT1) was isolated in Vero E6 cells from a nasopharyngeal specimen collected in April 2020. The recombinant fluorescent reporter virus icSARS-CoV-2-mNG was generated as described previously (6). The spike gene of all working stocks was sequenced. While USA-WA1 and MA/VPT1 did not have mutations or variations (at 20% cut off level), icSARS-CoV-2-mNG acquired a 5-residue insertion at the furin cleavage site resulting in a sequence change from “PRRARS” to “PRRNIGERARS” in majority of the viral population.

### Cells

MDCK-Atlanta, MDCK-London, and MDCK-SIAT1 cells were obtained from the International Reagent Resources (IRR). MDCK-hCK cells were kindly provided by Y. Kawaoka (University of Wisconsin-Madison). MDCK-NBL2, Vero E6, CV-1, A549, CRFK, Mv1Lu, RD, Hep-2c, HeLa, and L20B cells were obtained from American Type Culture Collection (ATCC) or maintained at CDC’s Division of Scientific Resources. Chicken embryo fibroblasts (CEF) were obtained from Charles River Laboratories (Wilmington, MA). All 25 cell lines listed in Table 1 were obtained from Quidel Corporation (San Diego, CA) in pre-seeded 24-well plates except for CRFK and RhMK cells, which were obtained in T-75 flasks and seeded into 24-well plates in the lab one day prior to infection.

**Table 1.** Overview of diagnostic cell lines obtained from a commercial source (Quidel).

| Cell line | Organism | Tissue | Type/<br>Morphology | Virus susceptibility profile* | SARS-CoV-1<br>susceptible | SARS-CoV-2<br>susceptible |
| --- | --- | --- | --- | --- | --- | --- |
| Vero | African green monkey | Kidney | Epithelial | AdV, coxsackie B, measles, mumps, rotavirus, rubella, influenza | Yes (28, 34) | Yes |
| Vero 76 | African green monkey | Kidney | Epithelial | AdV, coxsackie B, measles, mumps, poliovirus, rotavirus, rubella, West Nile Virus | Yes (35) | Yes |
| BGMK | African green monkey | Kidney | Epithelial | coxsackie B, poliovirus | Yes (28) | Yes |
| CV-1 | African green monkey | Kidney | Fibroblast | measles, mumps, rotavirus | Yes (28) | No |
| LLC-MK2 | Rhesus macaque | Kidney | Epithelial | enterovirus, myxovirus and poxvirus groups, poliovirus type 1, rhinovirus | Yes (28) | Yes |
| RhMK | Rhesus macaque | Kidney | Epithelial | enteroviruses, influenza, parainfluenza | Yes (31) | Yes |
| A549 | Human | Lung | Epithelial | AdV, influenza, measles, mumps, parainfluenza, poliovirus, RSV, rotavirus | No (28, 30, 31); Yes(36) | No |
| HEL | Human | Lung | Fibroblast | AdV, CMV, echovirus, HSV, poliovirus, rhinovirus | No (28, 31) | No |
| HeLa | Human | Cervix | Epithelial | AdV, CMV, echovirus, HSV, poliovirus, rhinovirus | No (28) | No |
| HeLa 229 | Human | Cervix | Epithelial | AdV, CMV, echovirus, HSV, poliovirus, rhinovirus | Unknown | No |
| HEp2 | Human | Cervix | Epithelial | AdV, coxsackie B, HSV, measles, parainfluenza, poliovirus, RSV | No (28) | No |
| MRC-5 | Human | Lung | Fibroblast | AdV, CMV, echovirus, HSV, influenza, mumps, poliovirus, rhinovirus | No (31) | No |
| MRHF | Human | Foreskin | Fibroblast | AdV, CMV, echovirus, HSV, mumps, poliovirus, rhinovirus | Unknown | No |
| NCI-H292 | Human | Lung | Epithelial | AdV, HSV, influenza A, measles virus, RSV, rhinoviruses, vaccinia virus | No (30, 33, 36) | No |
| RD | Human | Muscle | Spindle;<br>multi-nucleated | AdV, echovirus, HSV, poliovirus | No (28, 32) | No |
| WI-38 | Human | Lung | Fibroblast | AdV, CMV, echovirus, HSV, influenza, mumps, poliovirus, rhinovirus, RSV | Unknown | No |
| McCoy | Mouse | Unknown | Fibroblast | HSV | Unknown | No |
| MNA | Mouse | Nerve | Neuroblastoma | Rabies | Unknown | No |
| MDCK | Dog | Kidney | Epithelial | AdV, coxsackie virus, influenza, reoviruses | No (25, 28, 29, 31, 33) | No |
| CRFK | Cat | Kidney | Epithelial | canine parvovirus, feline calicivirus, feline panleukopenia virus, rabies virus | Yes (25) | Yes (limited) |
| Mv1Lu | American mink | Lung | Epithelial | CMV, influenza | Yes (31, 34) | No |
| H&V-Mix | CV-1 and MRC-5 | Mixture | Mixture | AdV, CMV, echovirus, HSV, influenza, poliovirus type 1, SV40 virus, VZV | Unknown | No |
| R-Mix | Mv1Lu and A549 | Mixture | Mixture | AdV, CMV, HSV, influenza, measles, mumps, poliovirus, RSV, rotavirus | Yes (31) | No |
| R-Mix Too | MDCK and A549 | Mixture | Mixture | AdV, HSV, influenza, MPV, measles, mumps, poliovirus, RSV, rotavirus, VZV | Unknown | No |
| Super E-Mix | BGMK and A549 | Mixture | Mixture | AdV, HSV, influenza, measles, mumps, poliovirus, RSV, rotavirus, VZV | Unknown | Yes |

\*Virus susceptibility profiles listed are as reported by Quidel and not verified in this study. AdV, adenovirus; CMV, cytomegalovirus; HSV, herpes simplex virus; RSV, respiratory syncytial virus; VZV, varicella zoster virus.

### Virus Infection of cell lines

Cells were seeded in 6-, 12-, or 24-well plates a day prior to infection or used directly upon receipt from a commercial source (Quidel). Infection dose for each experiment is specified in the results section or figure legends. Infection temperature was always 37°C. In general, inoculum was saved for back titration and the result was shown as 0 hours post inoculation (hpi) in some figures. Cells were then washed at 1-2 hpi and supernatants or cell lysates were collected daily for at least 3 days and up to 5 days for infectious virus titration and for viral RNA quantification, respectively. Cytopathic effect (CPE) and fluorescence signals (for icSARS-CoV-2-mNG) were observed daily.

### Virus infection of embryonated chicken eggs

Specific-pathogen-free (SPF) embryonated chicken eggs were obtained from Charles River Laboratories (North Franklin, CT, USA). USA-WA1 was inoculated into the allantoic cavity of 24 8-to 12-day-old eggs at 10^5^ TCID_50_/egg and incubated at 37°C for 3 days. Allantoic fluid was collected from individual eggs separately as E1 samples. One hundred μl of each E1 sample was passaged into a corresponding egg and 24 E2 samples were collected after 3 days of incubation. Similarly, 24 E3 samples were generated from passage of E2 samples in 24 eggs. All E1, E2, and E3 samples, as well as samples from cell lines, were titrated by TCID_50_ assay and viral RNAs were quantified by real-time reverse transcription PCR (rRT-PCR) assay (7). Synthetic RNA was used in the rRT-PCR assay to generate the standard curve for absolute quantification.

### Immunoblot detection of ACE2

Cells were lysed in NP-40 lysis buffer and protein concentrations were determined using a BCA protein assay kit (Pierce). Cell lysates and recombinant ACE2 protein control (Sino Biological) were immunoblotted for ACE2 and β-actin using primary antibodies (1:500 polyclonal goat anti-human ACE2, R&D Systems, AF933; 1:1000 monoclonal mouse anti-β-Actin, Abcam, AB8226) followed by secondary antibodies (1:4000 donkey anti-goat, Abcam; 1:4000 goat anti-mouse, Biorad). Immunoblots were developed using SuperSignal West Pico PLUS Chemiluminescent Substrate (ThermoFisher).

### Expression of recombinant ACE2 proteins

The Expi293 Expression system (ThermoFisher) was used for production of histidine-tagged ACE2 (ectodomain) proteins. The Expi293F cells were transfected with pCAGGS-ACE2 mammalian expression construct and cultured at 37°C with 8% CO2 at a shaking speed of 125 RPM. The supernatant was harvested on day 5 and ACE2 protein was purified using HisTrap FF column (GE Life Sciences), followed by desalting. The purified protein was further concentrated on Amicon Ultra Centrifugal Filters with 50 KDa cutoff (Sigma-Aldrich).

### Bio-layer interferometry assay

Affinity between SARS-CoV-2 S1 (Sino Biological, 40591-V02H) and human ACE2 (hACE2) or canine ACE2 (cACE2) were evaluated using Octet RED96 instrument at 30°C with a shaking speed of 1000 RPM (ForteBio). Anti-penta-His biosensors (HIS1K) (ForteBio) were used. hACE2 or cACE2 was loaded onto surface of biosensor at 100 nM in 10X kinetic buffer (ForteBio) for 5 minutes. After 1 minute of baseline equilibration, 5 minutes of association was conducted with 10-100 nM of S1 to hACE2 or 25-200 nM of S1 to cACE2, followed by 5 minutes of dissociation. The data were corrected by subtracting reference sample, and 1:2 (Bivalent) binding model with global fit was used for determination of affinity constants.

### Exogenous expression of ACE2 in MDCK cells

Constructs co-expressing full-length hACE2 or cACE2 with mCherry2 protein (CMV promoter-ACE2-IRES-mCherry2) were generated and transfected into MDCK-SIAT1 cells via electroporation with Lonza Nucleofector system (Lonza) using the manufacturer’s protocol with program A024. 1.5×10^6^ MDCK-SIAT1 cells were transfected with 10 μg DNA (pCMV-hACE2-IRES-mCherry2, pCMV-cACE2-IRES-mCherry2, or pCMV-IRES-mCherry2 empty control). One day post transfection, the cells were inoculated with USA-WA1 or icSARS-CoV-2-mNG.

### ACE2 Sequence alignment

ACE2 protein sequences for human (NP_001358344.1), African green monkey (AAY57872.1), rhesus macaque (ACI04564.1), mouse (NP_001123985.1), dog (XP_005641049.1), cat (NP_001034545.1), American mink (QPL12211), and chicken (XP_416822.2) were aligned using MUSCLE alignment in Geneious Prime software (version 2019.2.3).

## RESULTS

### Replication of SARS-CoV-2 in a large set of cell substrates from a commercial source

As the prevalence of SARS-CoV-2 infection increases during the pandemic, or when social-distancing restriction is relaxed in the post-pandemic era, additional coinfections with various human viruses are inevitable. Therefore, we assessed 25 cell substrates commercially available from Quidel (Table 1), some of which are widely used for virus diagnosis in clinical laboratories. The cells were seeded in 24-well plates and inoculated with 5×10^4^ TCID_50_/well of a fluorescent reporter virus in which the ORF7a gene was replaced by the mNeonGreen gene (icSARS-CoV-2-mNG), allowing successful infection to be visualized by green fluorescence signal (6). Almost all non-human primate cell lines in this panel were susceptible to icSARS-CoV-2-mNG infection except for CV-1 cells (Figure 1). In contrast, none of the human, mouse, mink, dog, or cat cell lines tested yielded fluorescent cells after infection. The Super-E Mix cells were likely susceptible because this cell culture is a mixture containing BGMK cells, which were found to be susceptible to SARS-CoV-2 (Figure 1). We then inoculated all these cell lines with 5×10^4^ TCID_50_/well of the wild type SARS-CoV-2/USA-WA1/2020 (USA-WA1) strain and titrated supernatants collected over 5 days. Consistent with the results from icSARS-CoV-2-mNG infection, all non-human primate cell lines except CV-1 cells supported productive virus replication, whereas all other cell lines failed to generate infectious virus (Figure 2). It should be noted that viral titers in CRFK cells increased slightly at 2 dpi (Figure 2), suggesting that this cell line may support a low level of replication. The results along with the cell substrates’ information are summarized in Table 1.

**Figure 1.**
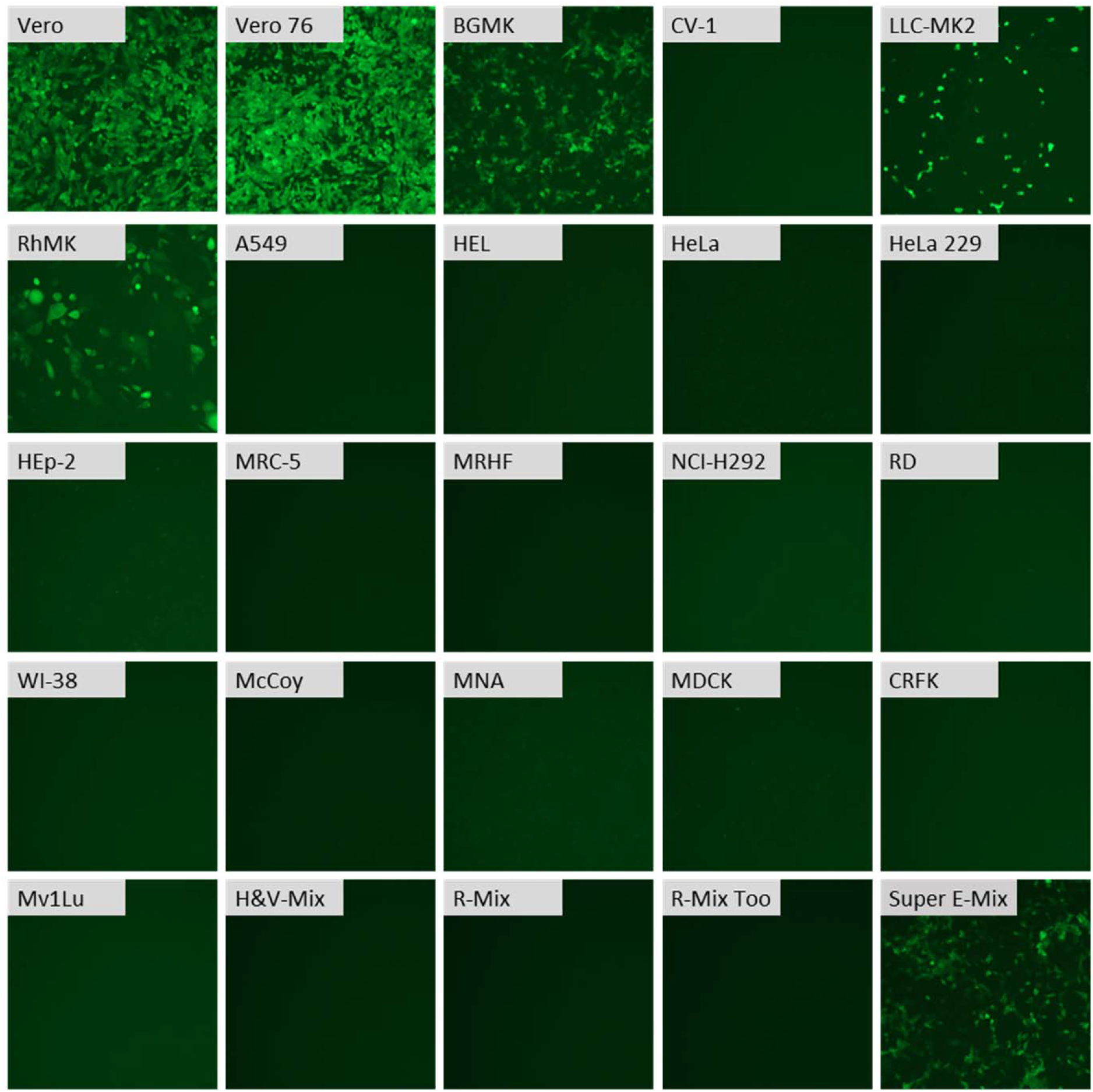
SARS-CoV-2 infects select commercially sourced cell lines. Cell lines were inoculated with the SARS-CoV-2 reporter virus encoding mNeonGreen (icSARS-CoV-2-mNG) at 5×10^4^ TCID_50_/well in 24-well plates, and infected cells were identified by green fluorescence. Microscopy images were captured at 24 hpi using 10X magnification. Representative images at 1 dpi are shown but similar results were observed through 5 dpi, and all mNeonGreen-negative cell lines remained negative.

**Figure 2.**
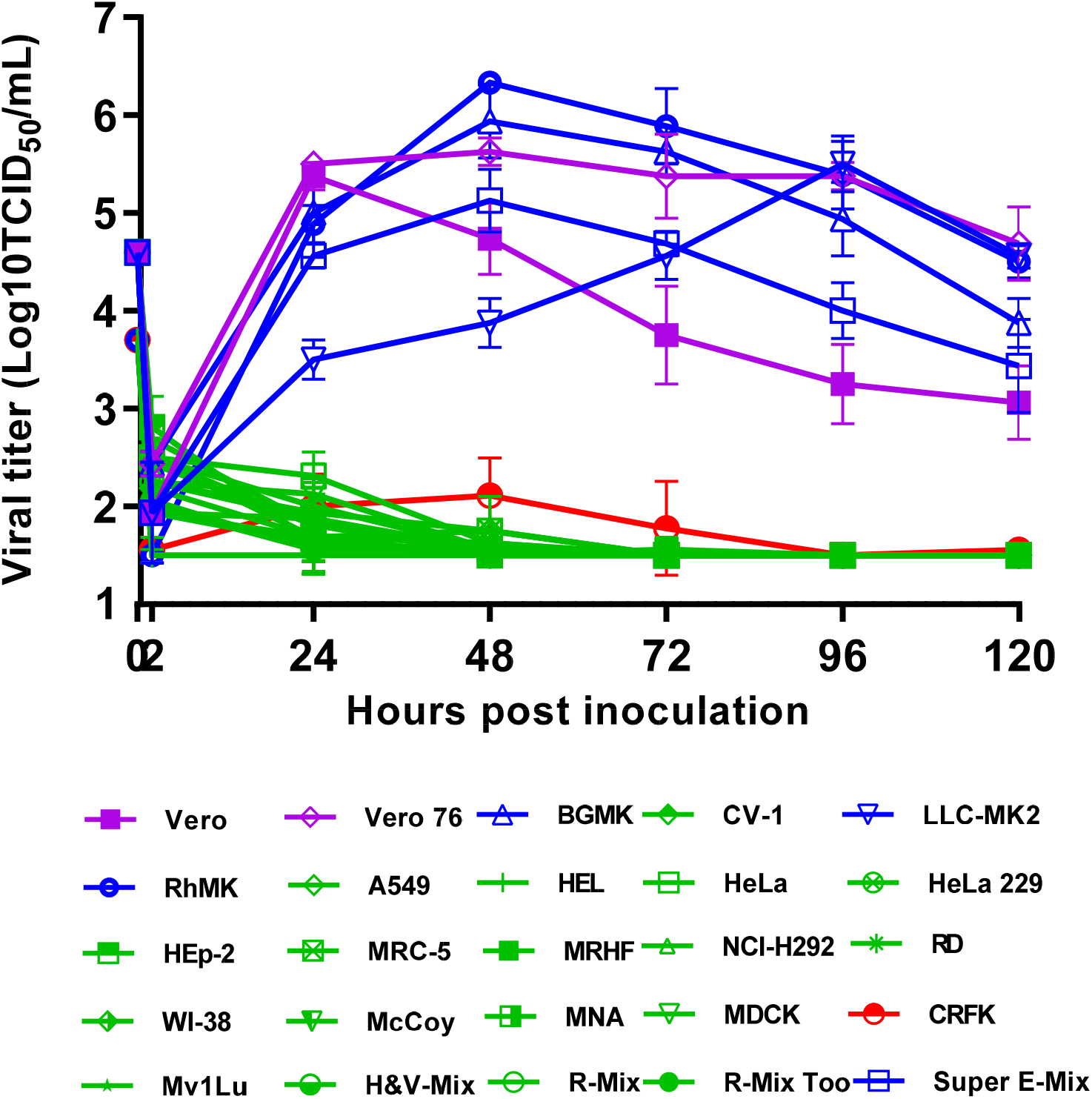
SARS-CoV-2 viral replication kinetics vary in commercially sourced cell lines. The 25 cell lines obtained from Quidel were inoculated with USA-WA1 at 5×10^4^ TCID_50_/well in 24-well plates, and supernatants were harvested at the indicated times and assayed for viral replication by TCID_50_ assay. Data are mean of n=4 ± sd.

### Replication of SARS-CoV-2 in influenza virus substrates

MDCK cells and embryonated chicken eggs are widely used for influenza virus isolation and propagation. There are multiple lineages or derivatives of MDCK cells used by laboratories for different types or subtypes of influenza viruses. Some lineages, such as MDCK-SIAT1 and hCK cells, were genetically modified and cloned from single cells, resulting in altered cell morphology and enhanced susceptibility to some subtypes of influenza viruses compared to their parental MDCK cell lines (8, 9). The different lineages of MDCK cells have altered gene expression profiles and surface glycans, and it is unclear whether that would affect their susceptibility to SARS-CoV-2. Therefore, we examined the susceptibility to SARS-CoV-2 in representative lineages of MDCK cells that are used widely in different laboratories, including MDCK-NBL-2, MDCK-Atlanta, MDCK-London, MDCK-SIAT1, and MDCK-hCK.

We inoculated Vero E6 cells (as a positive control) and various MDCK cell lines with 5×10^4^ TCID_50_/well of USA-WA1 and incubated for 1-2 hours at 37°C. Cells were then washed to remove the inoculum, and influenza virus infection media containing TPCK-trypsin and bovine serum albumin (BSA) was added to mimic the conditions used in influenza virus isolation. Supernatants were collected at the indicated times post-infection and viral titers measured. Vero E6 cells supported robust viral replication and reached peak titer within 2 days (Figure 3A), and infection killed most cells (data not shown). In contrast, none of the five MDCK cell lines tested supported SARS-CoV-2 replication. While residual infectious virus was present in some MDCK supernatant samples at 2 hpi, it was below the limit of detection (LOD) by 1-day post-infection (dpi) and did not cause any cytopathic effect (CPE) through 5 dpi. Similar experiments were conducted with the MDCK cell lines in which the infection media contained FBS rather than BSA, and again SARS-CoV-2 failed to replicate in any of the 5 MDCK cell lines (data not shown but almost identical to Figure 3A).

**Figure 3.**
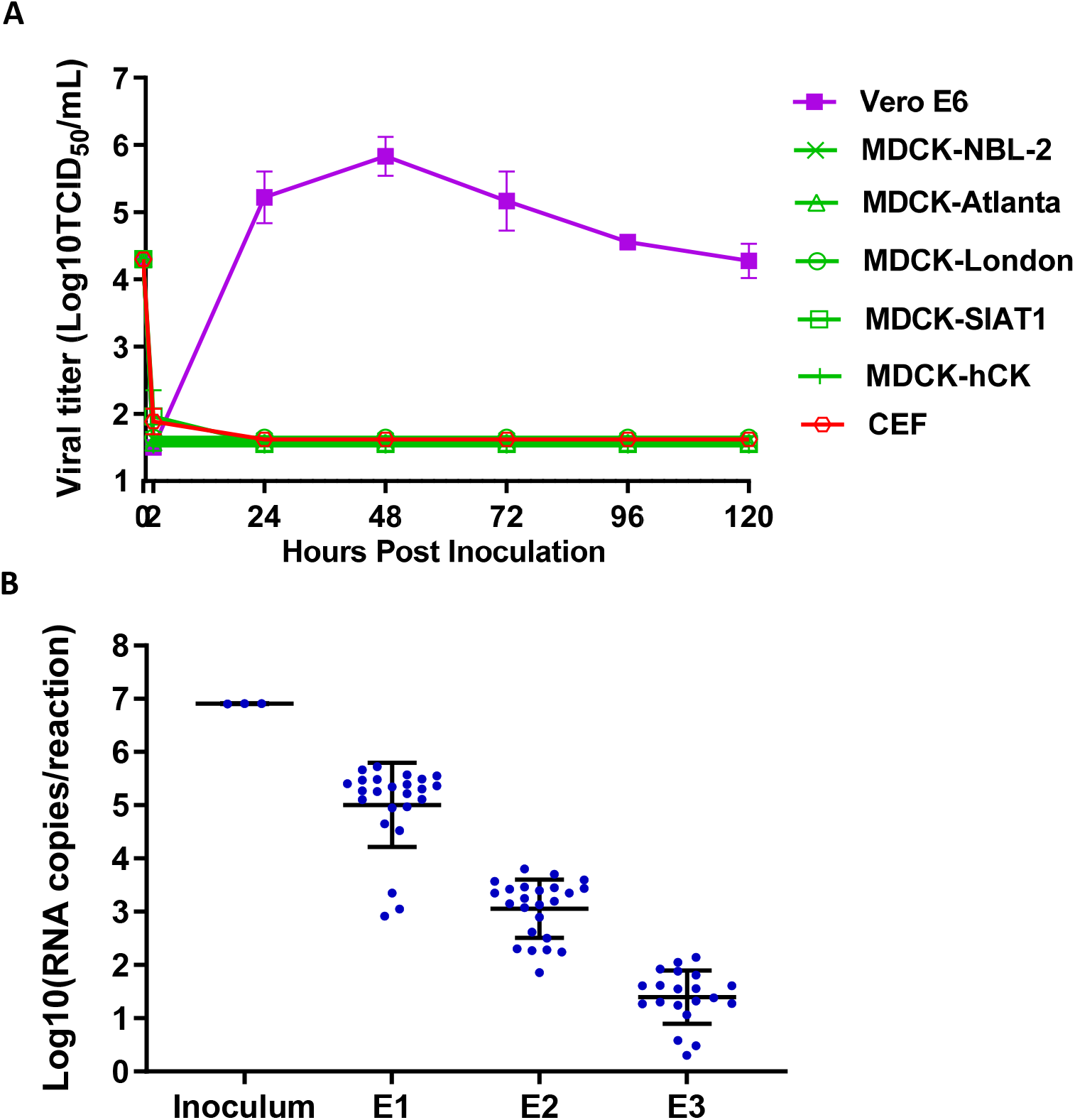
Influenza virus substrates do not support SARS-CoV-2 infection. (A) Vero E6, MDCK-NBL-2, MDCK-Atlanta, MDCK-London, MDCK-SIAT1, MDCK-hCK, and CEF cells were inoculated with USA-WA1 at 5×10^4^ TCID_50_/well in 12-well plates, and supernatant were harvested and assayed for viral replication by TCID_50_ assay. (B) USA-WA1 total viral RNA levels in allantoic fluid from infected eggs were quantified by rRT-PCR using a standard curve generated by synthetic RNA. Not plotted are four eggs with undetectable RNAs for E3. Data are a mean of n=3 ± sd (cells) or n=24 ± sd (eggs).

Embryonated chicken eggs are another common substrate for influenza virus isolation, propagation, and vaccine production. We inoculated 24 eggs each with 10^5^ TCID_50_ of USA-WA1 and blindly passaged the virus in eggs for 3 passages (E1, E2, and E3). Viral titers in the allantoic fluid of E1, E2, and E3 eggs were below the limit of detection (10^1.5^ TCID_50_/ml) even in E1 eggs (data not shown). We then used an rRT-PCR assay to quantify the viral RNA levels in the inoculum and allantoic fluid samples (7). Viral RNA decreased steadily over the 3 passages in eggs (Figure 3B). We also inoculated chicken embryo fibroblasts (CEF) with USA-WA1, and no infectious virus was produced from the cells (Figure 3A). These results clearly demonstrate that embryonated chicken eggs are not a susceptible substrate for the SARS-CoV-2 replication.

Collectively, the data show that substrates commonly used to culture influenza A and B viruses are not susceptible to SARS-CoV-2 infection.

### Replication of SARS-CoV-2 in polio and enterovirus substrates

From patients potentially infected with polio or enteroviruses, stool specimens are used to inoculate appropriate cell lines for surveillance. As SARS-CoV-2 virus can infect multiple organs and tissues and its presence in stool specimens has been reported (10–16), it is important to determine if cell lines commonly used for polio and enterovirus culture could inadvertently propagate SARS-CoV-2. Therefore, RD, HeLa, Hep-2C, and L20B cells were inoculated with USA-WA1 at MOI of 0.1 and incubated for 2 hours after which the inoculum was removed and cells were washed 3 times to remove residual virus. No CPE was observed over a 4-day period, and SARS-CoV-2 was not detectable in supernatant collected at 1-4 dpi (data not shown). This result was confirmed by rRT-PCR of cell lysate, which revealed that the total viral RNA levels decreased compared to the inoculum, indicating that virus did not efficiently initiate RNA transcription or replication (Figure 4). These results indicate that cell substrates regularly used for polio and enterovirus cultures are not susceptible to SARS-CoV-2 infection when cultured under these standard conditions.

**Figure 4.**
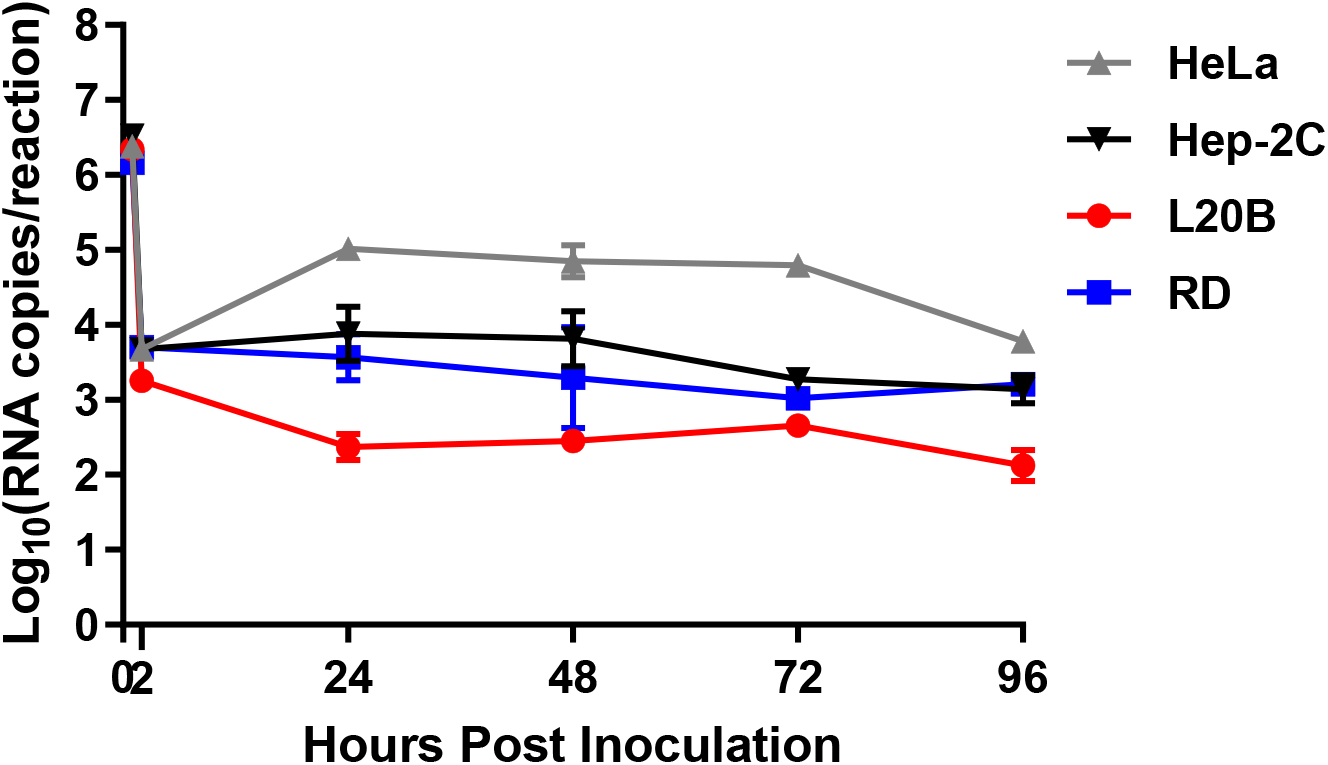
Poliovirus substrates do not support SARS-CoV-2 infection. Total viral RNA levels were determined by rRT-PCR (standard curve generated by synthetic RNA) from total RNA extracted from cell lines inoculated with USA-WA1 at MOI of 0.1 in 6-well plates. The data points at 1h are represented by the RNA from the inoculum while 2h and later time points are from RNA extracted from cell lysates. Data are mean of n=3 ± sd.

### Replication of SARS-CoV-2 with Spike-D614G substitution

During this study, we noticed that the proportion of naturally circulating virus containing a D614G substitution in the spike protein was rapidly increasing. The USA-WA1 strain is an early isolate that expresses spike with D614. To confirm that the cell susceptibility data obtained using this virus were valid with recent strains, a subset of representative cell lines were inoculated with high titer (5×10^5^ TCID_50_/well) of SARS-CoV-2/Massachusetts/VPT1/2020 (MA/VPT1), which encodes a spike with G614. In selection of cell lines for the subset, we included Vero E6 cells as a cell line that should support replication of MA/VPT1 given our previous findings with USA-WA1 (Figure 3A). Indeed, Vero E6 cells supported similar replication kinetics for MA/VPT1 as USA-WA1 (Figure 5A). Even with a 10-fold higher inoculum of MA/VPT1 than previously used for USA-WA1 tests (5×10^4^ TCID_50_/well), CV-1, A549, Mv1Lu, MDCK-NBL-2, and MDCK-SIAT1 cell lines were not susceptible to this SARS-CoV-2 strain encoding spike-G614. CRFK cells inoculated with MA/VPT1 generated virus titers slightly above the LOD at 1 dpi, after which titers decreased (Figure 5A). Viral titers were further confirmed by rRT-PCR. Similar to the virus titer data, inoculated CRFK cells had a 5-fold increase of viral RNA at 1 dpi compared to 2 hpi, but the RNA levels decreased over the next two days. In contrast, CV-1, A549, Mv1Lu, MDCK-NBL-2, and MDCK-SIAT1 cells did not shown any noticeable increase of viral RNA levels during the time course of this study (Figure 5B). All the 7 cell lines in this subset demonstrated very similar viral replication kinetics for both MA/VPT1 and USA-WA1 virus strains (Figures 2–5), indicating that the currently dominant virus strains with spike-G614 likely have the same cell susceptibility profile as earlier strains encoding spike-D614.

**Figure 5.**
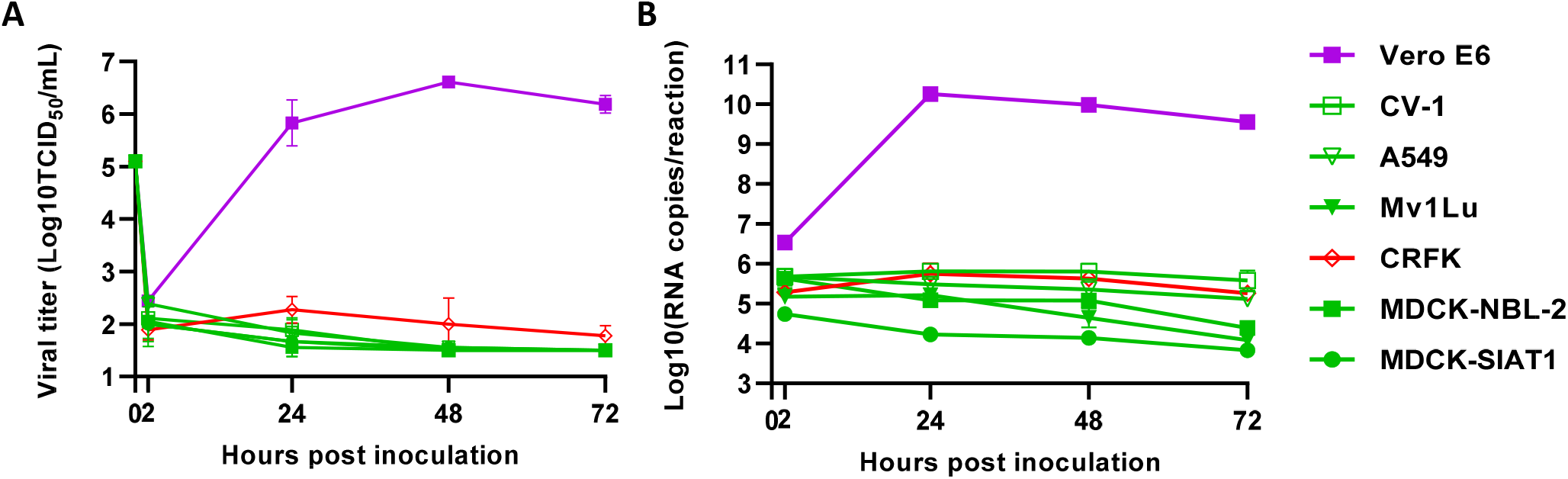
SARS-CoV-2 with spike-G614 infects similar cell types as SARS-CoV-2 with spike-D614. Vero E6, CV-1, A549, Mv1Lu, CRFK, MDCK-NBL-2, and MDCK-SIAT1 cell lines were inoculated with MA/VPT1 at 5×10^5^ TCID_50_/well in 12-well plates. (A) Supernatants were collected at the indicated times and viral replication kinetics determined using TCID_50_. Total RNA was extracted from cells inoculated for the indicated times and (B) total viral RNA levels were determined using rRT-PCR (standard curve generated by synthetic RNA). For all, data are a mean of n=3 ± sd.

### ACE2 as a critical determinant in susceptibility and species specificity

Coronavirus spike-host receptor interactions play the major role in species specificity (17). SARS-CoV-2 uses human angiotensin converting enzyme 2 (hACE2) as the host cell receptor (18). Multiple species including humans, monkeys, cats, minks, ferrets, hamsters, and dogs have been infected by SARS-CoV-2 in experimental and/or natural settings (19–24). To further investigate the mechanism of susceptibility or resistance and gain insight into SARS-CoV-2 species specificity, we analyzed the ACE2 expression levels in various cell lines. Multiple anti-ACE2 antibodies were screened to identify a polyclonal antibody that reacts with ACE2 from African Green Monkey (Vero and CV-1), Mink (Mv1Lu), Canine (MDCK), and feline (CRFK) (data not shown). Using this antibody, we determined by immunoblot that endogenous ACE2 levels were very high in Vero E6 cells derived from African Green Monkey kidney but extremely low in the other African green monkey kidney cell line CV-1, which could explain the drastic difference in infectivity between these two cell lines. Canine ACE2 protein was not detectable in MDCK cells, which surely plays a role in their resistance to SARS-CoV-2 infection. Similarly, the feline CRFK, mink Mv1Lu and human A549 cells had very low or undetectable endogenous ACE2 expression (Figure 6). The low protein levels of ACE2 in those cells coincided with low mRNA levels determined by rRT-PCR (data not shown).

**Figure 6.**
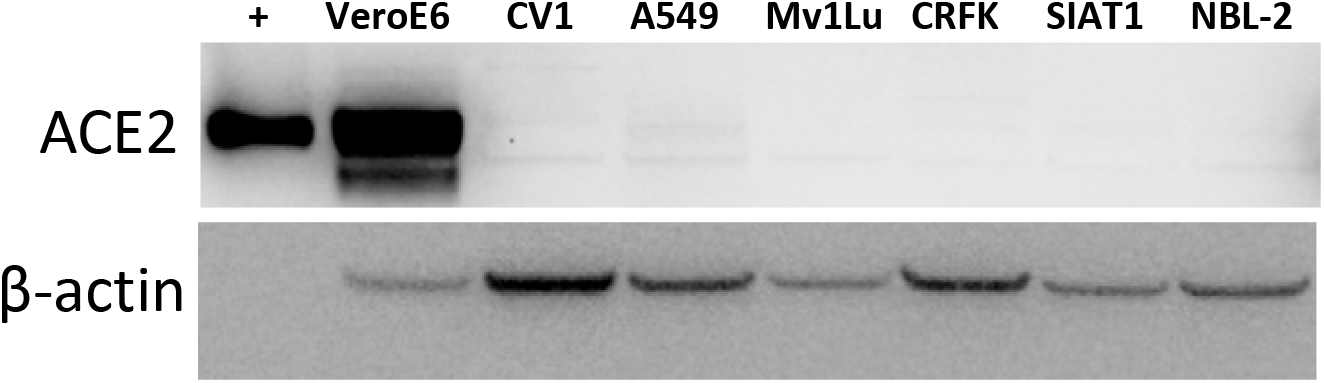
ACE2 is differentially expressed across cell lines. Whole cell lysate from uninoculated Vero E6, CV-1, A549, Mv1Lu, CRFK, MDCK-NBL-2 and MDCK-SIAT1 cell lines were immunoblotted for endogenous ACE2 expression. Recombinant hACE2 (Sino Biological) was used as a positive control for detection of hACE2. 20 μg of cell lysates or 0.2 ng of recombinant hACE2 protein were loaded. β-actin was also immunoblotted from samples as a loading control.

Since MDCK cells are the most important cell line for influenza virus isolation and propagation and dogs have been infected with SARS-CoV-2, we selected canine ACE2 (cACE2) for additional analysis. To better understand resistance of MDCK cells to SARS-CoV-2, constructs co-expressing ACE2 protein (hACE2 or cACE2) under a CMV promoter and mCherry2 protein through an IRES element were transfected into MDCK-SIAT1 cells. MDCK cells expressing hACE2 (MDCK-hACE2) or cACE2 (MDCK-cACE2) as determined by mCherry2 expression were efficiently infected by icSARS-CoV-2-mNG (Figure 7A). As a control, MDCK cells were also transfected with an empty vector plasmid that expresses mCherry2 via the IRES element but does not encode an ACE2 protein (MDCK-vector). Consistent with wild type MDCK cells the MDCK-vector control cells were not susceptible to SARS-CoV-2 (Figure 7A). These results were further confirmed by infecting MDCK-hACE2 and MDCK-cACE2 cells with the wild type virus USA-WA1 and assaying viral replication kinetics. Viral infectious titers and viral RNA levels were elevated in MDCK cells overexpressing either hACE2 or cACE2 relative to MDCK-vector cells (Figure 7B and 7C).

**Figure 7.**
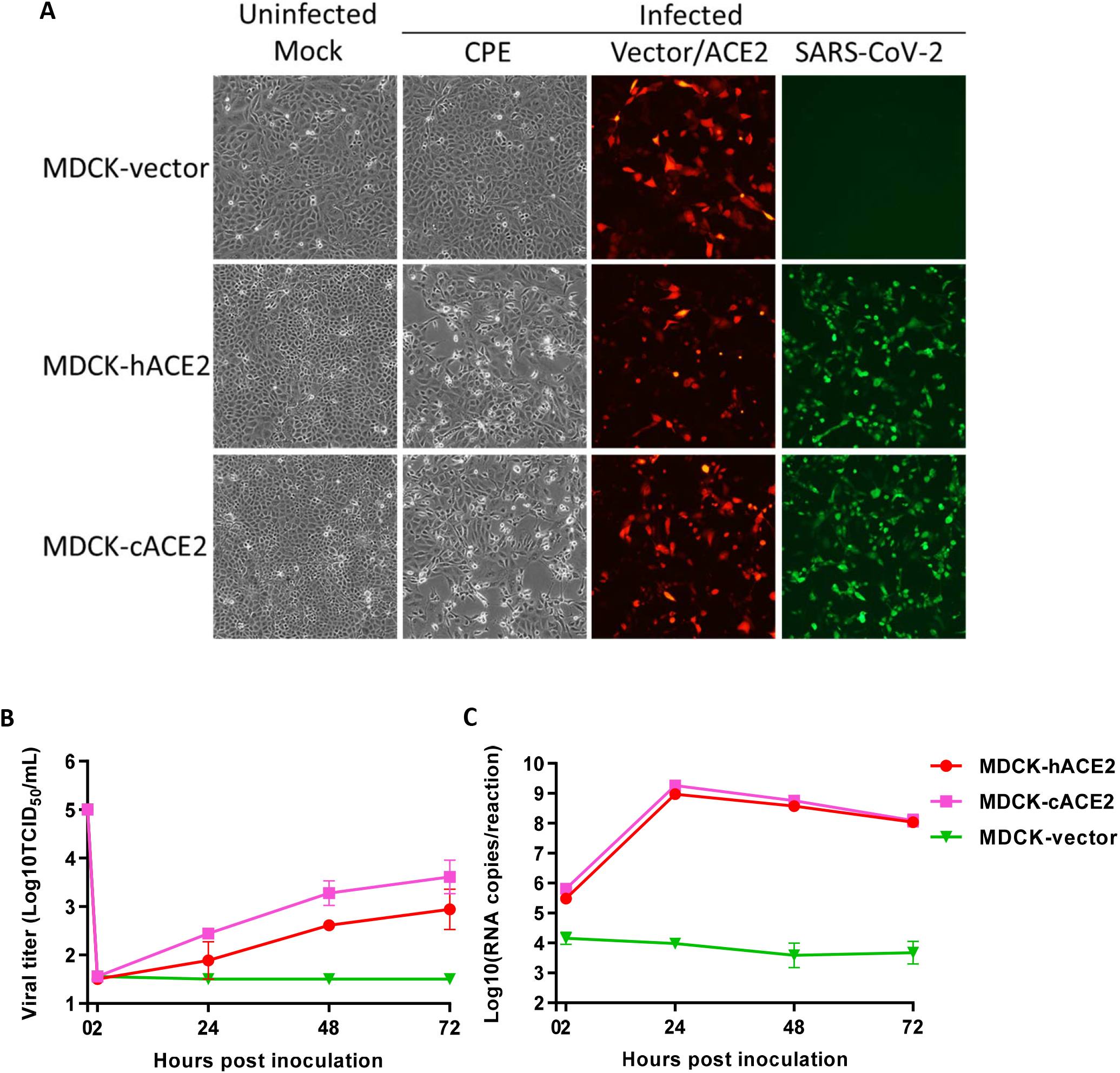
Overexpression of cACE2 permits SARS-CoV-2 infection of MDCK cells. (A) MDCK cells transiently overexpressing an empty vector control (MDCK-vector), hACE2 (MDCK-hACE2) or cACE2 (MDCK-cACE2) were mock-inoculated or inoculated with icSARS-CoV-2-mNG reporter virus at 5×10^5^ TCID_50_/well in 12-well plates, and viral infection detected by fluorescent microscopy. CPE was also imaged in inoculated cells. Representative images at 1 dpi are shown. (B-C) MDCK-vector, MDCK-hACE2, and MDCK-cACE2 cells were inoculated with USA-WA1 at 5×10^5^ TCID_50_/well in 12-well plates. Supernatants were collected at the indicated times and (B) viral titers determined by TCID_50_ assay. Total RNA was extracted from cell lines inoculated for the indicated length of time and (C) total viral RNA were determined using rRT-PCR (standard curve generated by synthetic RNA). Data for (B-C) are a mean of n=3 ± sd.

These results indicate that the resistance of MDCK cells to SARS-CoV-2 occurs at the virus entry step. Once bound, the genome is released, transcribed, translated, replicated and packaged into particles that bud from infected cells fairly efficiently. However, overexpression of ACE2 in MDCK cells could result in greater ACE2 expression as compared to most natural cell lines. Thus, even if cACE2 does not bind the spike protein as efficiently as hACE2, overexpression could facilitate entry of SARS-CoV-2 into MDCK-cACE2 cells. To determine if cACE2 binding affinity to SARS-CoV-2 spike was an additional factor preventing infection of MDCK cells, we conducted bio-layer interferometry (BLI) assays to compare the binding affinity of spike with cACE2 and hACE2. We identified that SARS-CoV-2 spike bound to cACE2 (KD = 19.5 nM) 15-fold less efficiently than hACE2 (KD = 1.30 nM) (Figure 8). The reduced binding affinity to cACE2 is likely a result of the sequence differences between the hACE and cACE2 in regions directly involved in spike binding (Figure 9). Thus, both low expression of cACE2 by MDCK cells and low binding affinity of cACE2 to SARS-CoV-2 spike contribute to the resistance of MDCK cells to SARS-CoV-2.

**Figure 8.**
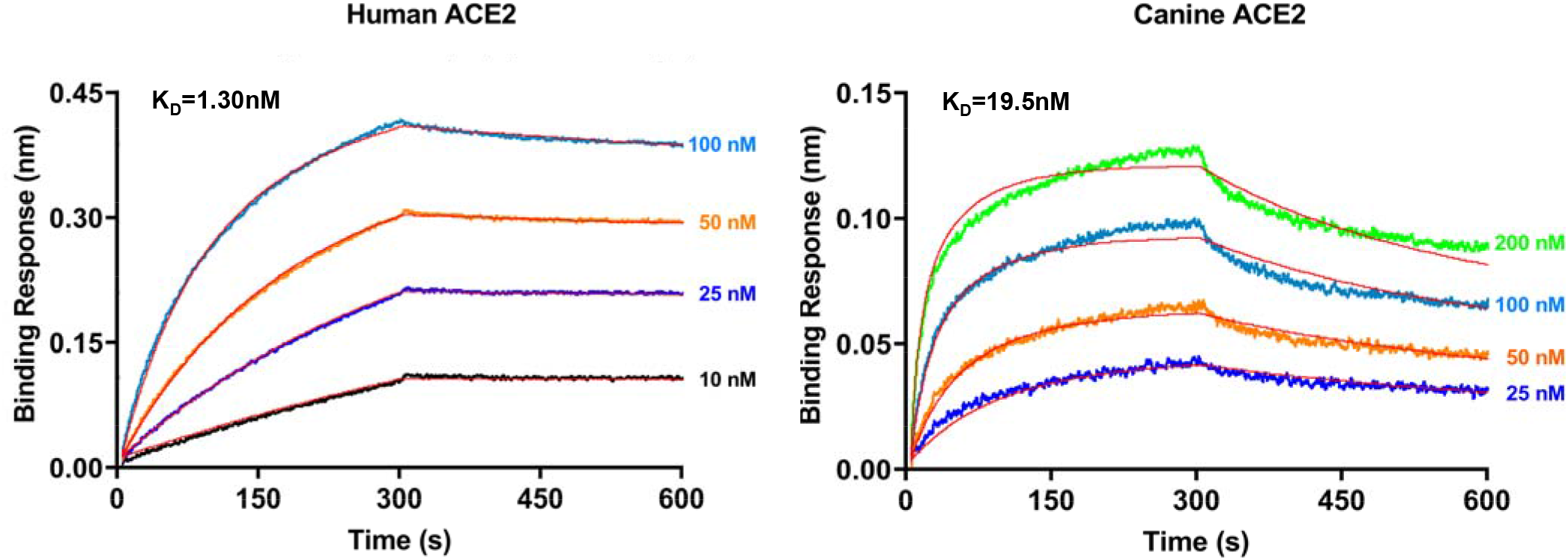
Canine ACE2 has lower affinity to SARS-CoV-2 spike protein compared to human ACE2. Bio-layer interferometry assay was used to determine the equilibrium dissociation constant (KD) of hACE2 or cACE2 protein with SARS-CoV-2 spike protein. hACE2 or cACE2 recombinant protein was loaded onto surface of biosensor at 100 nM and association was conducted at 10-100 nM of S1-Fc for hACE2 and 25-200 nM of S1-Fc for cACE2, followed by dissociation.

**Figure 9.**
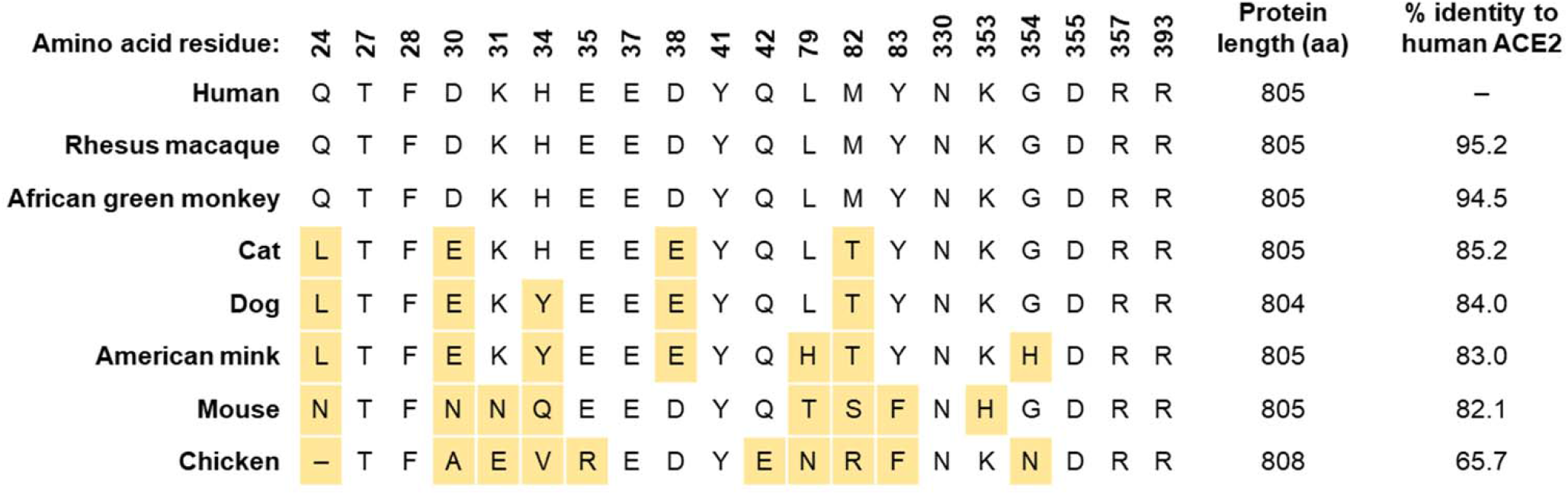
ACE2 protein sequences vary across species. ACE2 protein sequences from human, rhesus macaque, African green monkey, cat, dog, American mink, mouse, and chicken were aligned using MUSCLE. Residues involved in interaction with SARS-CoV-2 spike protein (based on ref (37–40)) are shown using hACE2 numbering, and residues varying from hACE2 are highlighted in yellow. A gap in alignment is indicated with a dash. Percent identity to hACE2 across the entire protein is shown.

## DISCUSSION

In this study, we determined the SARS-CoV-2 susceptibility of more than 30 cell lines or derivatives and embryonated chicken eggs. This study corroborates and complements other susceptibility studies published in the past few months (25, 26). For example, Barr *et al.* recently showed that MDCK cells and embryonated eggs do not support productive SARS-CoV-2 infection (26). The data presented here are consistent with that study, and our infectious virus titration assay data further showed that SARS-CoV-2 loses infectivity rapidly in cells and eggs, while the viral RNA levels decreased quite slowly. In addition, the majority of currently circulating strains contain the D614G substitution in the spike protein, which could impact binding, entry, and/or species specificity, and viruses with this change were not tested in previous studies. Herein, we showed that the spike-D614G substitution does not alter cell susceptibility of the cell lines tested including those with low levels of human (A549), non-human primate (CV-1), mink (Mv1Lu), cat (CRFK), or dog (MDCK) ACE2. In the future, even in the unlikely event that other spike substitutions render the binding of spike to cACE2 stronger (Fig 8), the low expression level of cACE2 in MDCK cells (Figure 6) still poses a high barrier for SARS-CoV-2 to overcome. Therefore, two independent studies together illustrate that MDCK cells and commonly utilized derivatives are not susceptible to SARS-CoV-2 and can be safely used for influenza virus isolation, propagation, and vaccine production. Additionally, chicken eggs which are used to manufacture most influenza virus vaccines do not support replication of SARS-CoV-2.

We expanded our examination to other clinically relevant cell lines used in diagnosis and isolation of a wide array of human viruses, particularly respiratory viruses (Table 1). While many of those cells were tested with SARS-CoV-1 virus previously (25, 27–36), it is worth noting that cell susceptibility conclusions derived from SARS-CoV-1 studies do not always apply to SARS-CoV-2. For example, we and others previously showed that Mv1Lu cells supported a moderate level of SARS-CoV-1 virus replication (31, 34), but they are not susceptible to SARS-CoV-2 replication as demonstrated in this study. This finding could be justified by the difference in ACE2 binding positions between SARS-CoV-1 and SARS-CoV-2 (37–40). Considering that mink ACE2 is only 83% identical to human ACE2 (Figure 9), some of the different ACE2 residues may have more adverse impact on the SARS-CoV-2 entry than on the SARS-CoV-1 entry. This idea does not necessarily contradict recent reports of SARS-CoV-2 infections among mink on farms (20, 41–44); ACE2 expression is relatively low in Mv1Lu cells (Figure 6) but likely higher in various epithelial cells *in vivo,* enabling productive infection in minks even through a weaker spike-receptor interaction.

Overall, our study provides important information on multiple cell lines and chicken eggs regarding their susceptibility to SARS-CoV-2. This study is important from a biosafety standpoint; humans can be coinfected by multiple pathogens. Specimens collected for testing and culture may contain SARS-CoV-2 and these data should help laboratories avoid inadvertent propagation. The data on canine ACE2 shed light on the relationship between SARS-CoV-2 susceptibility and ACE2 receptor affinity (species specificity) and expression level, suggesting that even ACE2 proteins with a number of substitutions at key residues that contact SARS-CoV-2 spike protein can still serve as functional receptors when expressed at high levels.

## Acknowledgements

We thank the support and guidance from the US Centers for Disease Control and Prevention COVID-19 Response Laboratory and Testing Task Force. We also thank the CDC Division of Scientific Resources for providing some cell lines and other materials. This activity was reviewed by CDC and was conducted consistent with applicable federal law and CDC policy: 45 C.F.R. part 46, 21 C.F.R. part 56; 42 U.S.C. Sect. 241(d); 5 U.S.C. Sect. 552a; 44 U.S.C. Sect. 3501 et seq. The conclusions, findings, and opinions expressed by authors contributing to this journal do not necessarily reflect the official position of the U.S. Department of Health and Human Services, the Public Health Service, the Centers for Disease Control and Prevention, or the authors’ affiliated institutions. Use of trade names is for identification only and does not imply endorsement by the Public Health Service or by the U.S. Department of Health and Human Services.

## References

1. Kim D, Quinn J, Pinsky B, Shah NH, Brown I. Rates of Co-infection Between SARS-CoV-2 and Other Respiratory Pathogens. Jama. 2020 Apr 15;323(20):2085–6.

2. Li ZT, Chen ZM, Chen LD, Zhan YQ, Li SQ, Cheng J, et al. Coinfection with SARS-CoV-2 and other respiratory pathogens in COVID-19 patients in Guangzhou, China. Journal of medical virology. 2020 May 28.

3. Konala VM, Adapa S, Naramala S, Chenna A, Lamichhane S, Garlapati PR, et al. A Case Series of Patients Coinfected With Influenza and COVID-19. Journal of investigative medicine high impact case reports. 2020 Jan-Dec;8:2324709620934674.

4. Yue H, Zhang M, Xing L, Wang K, Rao X, Liu H, et al. The epidemiology and clinical characteristics of co-infection of SARS-CoV-2 and influenza viruses in patients during COVID-19 outbreak. Journal of medical virology. 2020 Jun 12.

5. Harcourt J, Tamin A, Lu X, Kamili S, Sakthivel SK, Murray J, et al. Severe Acute Respiratory Syndrome Coronavirus 2 from Patient with Coronavirus Disease, United States. Emerg Infect Dis. 2020;26(6):1266–73.

6. Xie X, Muruato A, Lokugamage KG, Narayanan K, Zhang X, Zou J, et al. An Infectious cDNA Clone of SARS-CoV-2. Cell Host Microbe. 2020;27(5):841–8.e3.

7. Lu X, Wang L, Sakthivel S, Whitaker B, Murray J, Kamili S, et al. US CDC Real-Time Reverse Transcription PCR Panel for Detection of Severe Acute Respiratory Syndrome Coronavirus 2. Emerging Infectious Disease journal. 2020;26(8):1654.

8. Matrosovich M, Matrosovich T, Carr J, Roberts NA, Klenk HD. Overexpression of the alpha-2,6-sialyltransferase in MDCK cells increases influenza virus sensitivity to neuraminidase inhibitors. J Virol. 2003 Aug;77(15):8418–25.

9. Takada K, Kawakami C, Fan S, Chiba S, Zhong G, Gu C, et al. A humanized MDCK cell line for the efficient isolation and propagation of human influenza viruses. Nat Microbiol. 2019 Aug;4(8):1268–73.

10. Wang W, Xu Y, Gao R, Lu R, Han K, Wu G, et al. Detection of SARS-CoV-2 in Different Types of Clinical Specimens. Jama. 2020 Mar 11;323(18):1843–4.

11. Cheung KS, Hung IFN, Chan PPY, Lung KC, Tso E, Liu R, et al. Gastrointestinal Manifestations of SARS-CoV-2 Infection and Virus Load in Fecal Samples From a Hong Kong Cohort: Systematic Review and Meta-analysis. Gastroenterology. 2020 Jul;159(1):81–95.

12. Young BE, Ong SWX, Kalimuddin S, Low JG, Tan SY, Loh J, et al. Epidemiologic Features and Clinical Course of Patients Infected With SARS-CoV-2 in Singapore. Jama. 2020 Mar 3;323(15):1488–94.

13. Xu Y, Li X, Zhu B, Liang H, Fang C, Gong Y, et al. Characteristics of pediatric SARS-CoV-2 infection and potential evidence for persistent fecal viral shedding. Nature medicine. 2020 Apr;26(4):502–5.

14. Zheng S, Fan J, Yu F, Feng B, Lou B, Zou Q, et al. Viral load dynamics and disease severity in patients infected with SARS-CoV-2 in Zhejiang province, China, January-March 2020: retrospective cohort study. BMJ (Clinical research ed). 2020 Apr 21;369:m1443.

15. Tang A, Tong ZD, Wang HL, Dai YX, Li KF, Liu JN, et al. Detection of Novel Coronavirus by RT-PCR in Stool Specimen from Asymptomatic Child, China. Emerg Infect Dis. 2020 Jun;26(6):1337–9.

16. Clinical and virologic characteristics of the first 12 patients with coronavirus disease 2019 (COVID-19) in the United States. Nature medicine. 2020 Jun;26(6):861–8.

17. wentworth d. Coronavirus Binding and Entry’. Coronaviruses: Molecular and Cellular Biology: caister academic press; 2007.

18. Zhang H, Penninger JM, Li Y, Zhong N, Slutsky AS. Angiotensin-converting enzyme 2 (ACE2) as a SARS-CoV-2 receptor: molecular mechanisms and potential therapeutic target. Intensive Care Med. 2020/03/04 ed; 2020. p. 586–90.

19. Abdel-Moneim AS, Abdelwhab EM. Evidence for SARS-CoV-2 Infection of Animal Hosts. Pathogens. 2020 Jun 30;9(7).

20. Oreshkova N, Molenaar RJ, Vreman S, Harders F, Oude Munnink BB, Hakze-van der Honing RW, et al. SARS-CoV-2 infection in farmed minks, the Netherlands, April and May 2020. Euro surveillance: bulletin Europeen sur les maladies transmissibles = European communicable disease bulletin. 2020 Jun;25(23).

21. Munster VJ, Feldmann F, Williamson BN, van Doremalen N, Perez-Perez L, Schulz J, et al. Respiratory disease in rhesus macaques inoculated with SARS-CoV-2. Nature. 2020 Sep;585(7824):268–72.

22. Bosco-Lauth AM, Hartwig AE, Porter SM, Gordy PW, Nehring M, Byas AD, et al. Experimental infection of domestic dogs and cats with SARS-CoV-2: Pathogenesis, transmission, and response to reexposure in cats. Proc Natl Acad Sci U S A. 2020 Oct 20;117(42):26382–8.

23. Halfmann PJ, Hatta M, Chiba S, Maemura T, Fan S, Takeda M, et al. Transmission of SARS-CoV-2 in Domestic Cats. N Engl J Med. 2020 Aug 6;383(6):592–4.

24. Singla R, Mishra A, Joshi R, Jha S, Sharma AR, Upadhyay S, et al. Human animal interface of SARS-CoV-2 (COVID-19) transmission: a critical appraisal of scientific evidence. Vet Res Commun. 2020 Nov;44(3-4):119–30.

25. Chu H, Chan JF, Yuen TT, Shuai H, Yuan S, Wang Y, et al. Comparative tropism, replication kinetics, and cell damage profiling of SARS-CoV-2 and SARS-CoV with implications for clinical manifestations, transmissibility, and laboratory studies of COVID-19: an observational study. Lancet Microbe. 2020 May;1(1):e14–e23.

26. Barr IG, Rynehart C, Whitney P, Druce J. SARS-CoV-2 does not replicate in embryonated hen’s eggs or in MDCK cell lines. 2020;25(25):2001122.

27. Hattermann K, Müller MA, Nitsche A, Wendt S, Donoso Mantke O, Niedrig M. Susceptibility of different eukaryotic cell lines to SARS-coronavirus. Arch Virol. 2005;150(5):1023–31.

28. Kaye M. SARS-associated coronavirus replication in cell lines. Emerg Infect Dis. 2006 Jan;12(1):128–33.

29. Yamashita M, Yamate M, Li GM, Ikuta K. Susceptibility of human and rat neural cell lines to infection by SARS-coronavirus. Biochem Biophys Res Commun. 2005 Aug 19;334(1):79–85.

30. Drosten C, Gunther S, Preiser W, van der Werf S, Brodt HR, Becker S, et al. Identification of a novel coronavirus in patients with severe acute respiratory syndrome. N Engl J Med. 2003 May 15;348(20):1967–76.

31. Gillim-Ross L, Taylor J, Scholl DR, Ridenour J, Masters PS, Wentworth DE. Discovery of novel human and animal cells infected by the severe acute respiratory syndrome coronavirus by replication-specific multiplex reverse transcription-PCR. J Clin Microbiol. 2004 Jul;42(7):3196–206.

32. Hattermann K, Muller MA, Nitsche A, Wendt S, Donoso Mantke O, Niedrig M. Susceptibility of different eukaryotic cell lines to SARS-coronavirus. Arch Virol. 2005 May;150(5):1023–31.

33. Ksiazek TG, Erdman D, Goldsmith CS, Zaki SR, Peret T, Emery S, et al. A novel coronavirus associated with severe acute respiratory syndrome. N Engl J Med. 2003 May 15;348(20):1953–66.

34. Mossel EC, Huang C, Narayanan K, Makino S, Tesh RB, Peters CJ. Exogenous ACE2 expression allows refractory cell lines to support severe acute respiratory syndrome coronavirus replication. J Virol. 2005 Mar;79(6):3846–50.

35. Severson WE, Shindo N, Sosa M, Fletcher T, 3rd, White EL, Ananthan S, et al. Development and validation of a high-throughput screen for inhibitors of SARS CoV and its application in screening of a 100,000-compound library. J Biomol Screen. 2007 Feb;12(1):33–40.

36. Yen YT, Liao F, Hsiao CH, Kao CL, Chen YC, Wu-Hsieh BA. Modeling the early events of severe acute respiratory syndrome coronavirus infection in vitro. J Virol. 2006 Mar;80(6):2684–93.

37. Yan R, Zhang Y, Li Y, Xia L, Guo Y, Zhou Q. Structural basis for the recognition of SARS-CoV-2 by full-length human ACE2. Science. 2020 Mar 27;367(6485):1444–8.

38. Wang Q, Zhang Y, Wu L, Niu S, Song C, Zhang Z, et al. Structural and Functional Basis of SARS-CoV-2 Entry by Using Human ACE2. Cell. 2020 May 14;181(4):894–904.e9.

39. Lan J, Ge J, Yu J, Shan S, Zhou H, Fan S, et al. Structure of the SARS-CoV-2 spike receptor-binding domain bound to the ACE2 receptor. Nature. 2020 May;581(7807):215–20.

40. Shang J, Ye G, Shi K, Wan Y, Luo C, Aihara H, et al. Structural basis of receptor recognition by SARS-CoV-2. Nature. 2020 May;581(7807):221–4.

41. Enserink M. Coronavirus rips through Dutch mink farms, triggering culls. Science. 2020 Jun 12;368(6496):1169.

42. Molenaar RJ, Vreman S, Hakze-van der Honing RW, Zwart R, de Rond J, Weesendorp E, et al. Clinical and Pathological Findings in SARS-CoV-2 Disease Outbreaks in Farmed Mink (Neovison vison). Veterinary pathology. 2020 Jul 14:300985820943535.

43. Hammer AS, Quaade ML, Rasmussen TB, Fonager J, Rasmussen M, Mundbjerg K, et al. SARS-CoV-2 Transmission between Mink (Neovison vison) and Humans, Denmark. Emerg Infect Dis. 2020 Nov 18;27(2).

44. Oude Munnink BB, Sikkema RS, Nieuwenhuijse DF, Molenaar RJ, Munger E, Molenkamp R, et al. Transmission of SARS-CoV-2 on mink farms between humans and mink and back to humans. Science. 2020 Nov 10.

